# Peroxisomes control mitochondrial dynamics and the mitochondrion-dependent pathway of apoptosis

**DOI:** 10.1101/407098

**Authors:** Hideaki Tanaka, Tomohiko Okazaki, Mutsumi Yokota, Masato Koike, Yasushi Okada, Yukio Fujiki, Yukiko Gotoh

**Affiliations:** Graduate School of Pharmaceutical Sciences, IRCN, The University of Tokyo, Tokyo 113-0033, Japan; Department of Cell Biology and Neuroscience, Juntendo University School of Medicine, Tokyo 113-8421, Japan; Laboratory for Cell Dynamics Observation, Center for Biosystems Dynamics Research (BDR), RIKEN, Osaka 565-0874, Japan; Department of Physics, Universal Biology Institute (UBI), and the International Research Center for Neurointelligence (WPI-IRCN), The University of Tokyo, Tokyo 113-0033, Japan; Division of Organelle Homeostasis, Medical Institute of Bioregulation, Kyushu University, Fukuoka 812-8582, Japan.

**Keywords:** organelle, peroxisome, mitochondria, apoptosis, Drp1, caspase

## Abstract

**Summary Statements:** We unveil a previously unrecognized role of peroxisomes in the regulation of mitochondrial fission-fusion dynamics, mitochondrion-dependent caspase activation, and cellular apoptosis.

**Abstract:** Peroxisomes cooperate with mitochondria in the performance of cellular metabolic functions such as fatty acid oxidation and maintenance of redox homeostasis. Whether peroxisomes also regulate mitochondrial fission-fusion dynamics or mitochondrion-dependent apoptosis has remained unclear, however. We now show that genetic ablation of the peroxins Pex3 or Pex5, which are essential for peroxisome biogenesis, resulted in mitochondrial fragmentation in mouse embryonic fibroblasts (MEFs) in a manner dependent on dynamin-related protein 1 (Drp1). Conversely, treatment with 4-phenylbutyric acid, an inducer of peroxisome proliferation, resulted in mitochondrial elongation in wild-type MEFs, but not in Pex3-deficient MEFs. We further found that peroxisome deficiency increased the levels of cytosolic cytochrome *c* and caspase activity under basal conditions without inducing apoptosis. It also greatly enhanced etoposide-induced caspase activation and apoptosis, indicative of an enhanced cellular sensitivity to death signals. Together, our data unveil a previously unrecognized role of peroxisomes in the regulation of mitochondrial dynamics and mitochondrion-dependent apoptosis. Given that mutations of peroxin genes are responsible for lethal disorders such as Zellweger syndrome, effects of such mutations on mitochondrion-dependent apoptosis may contribute to disease pathogenesis.

## Introduction

Peroxisomes are organelles bound by a single membrane that play essential roles in metabolic functions such as oxidation of fatty acid chains, catabolism of reactive oxygen species (ROS), and synthesis of ether phospholipids in all eukaryotic cells. The peroxin (Pex) family of proteins is required for the assembly and function of peroxisomes (Fujiki et al., 2012; Waterham and Ebberink, 2012). Deficiency of Pex3, a peroxisomal membrane protein necessary for membrane assembly, thus results in the complete loss of peroxisomes (Muntau et al., 2000), whereas deficiency of Pex5, a peroxisomal transporter, results in the loss of peroxisomal matrix proteins (Otera et al., 1998). Peroxisomal dysfunction due to *Pex* gene mutations is detrimental to human development, as evidenced by human autosomal recessive genetic diseases (known as peroxisome biogenesis disorders) such as Zellweger syndrome that result in death within the 1st year of life (Goldfischer et al., 1973). Peroxisome-deficient mice also die during the neonatal period (Baes et al., 1997; Maxwell et al., 2003). Both patients with peroxisome biogenesis disorders and peroxisome-deficient mice manifest a variety of characteristics including neurological dysfunction, hypotonia, and craniofacial abnormalities (Muntau et al., 2000; Trompier et al., 2014; Waterham and Ebberink, 2012).

Peroxisomes collaborate with other organelles in various physiological and pathological contexts. In particular, peroxisomes engage in a functional interplay with mitochondria with regard to the degradation of fatty acids and ROS detoxification as well as to antiviral immunity (Dixit et al., 2010; Lismont et al., 2015; Schrader and Yoon, 2007). The interplay between peroxisomes and mitochondria is highlighted by the observation that the loss of *Pex* genes gives rise to abnormalities in mitochondrial structure and metabolic function. For instance, deletion of *Pex5* in mouse hepatocytes affects the structure of mitochondrial inner and outer membranes as well as gives rise to abnormal (swollen) cristae (Baumgart et al., 2001; Goldfischer et al., 1973; Peeters et al., 2015), reduced activity of oxidative phosphorylation (OXPHOS) complexes, loss of the mitochondrial membrane potential, and increased ROS levels (Peeters et al., 2015). Deletion of *Pex5* was also shown to increase the number of mitochondria as well as the level of glycolytic activity, possibly as a compensatory response to the impairment of OXPHOS (Peeters et al., 2015). Deletion of *Pex13* in mouse brain or of *Pex19* in fly larvae resulted in similar dysfunction of OXPHOS, elevated ROS levels, and an increased abundance of mitochondria (Bülow et al., 2018; Rahim et al., 2016). Furthermore, human patients harboring *PEX16* mutations manifest myopathy accompanied by mitochondrial abnormalities (Salpietro et al., 2015). It remains unclear, however, which of these various phenotypes in mice, flies, and humans reflect primary effects of peroxisome deficiency or are secondary to primary effects such as ROS accumulation.

In addition to their roles in OXPHOS and redox regulation, mitochondria are key players in the regulation of apoptosis. Various proteins that are normally localized to the intermembrane space of mitochondria, including cytochrome *c*, are released into the cytosol on apoptosis induction. In the cytosol, cytochrome *c* interacts with Apaf-1 and pro-caspase-9 to form a large protein complex known as the apoptosome. The resulting increase in the autocatalytic activity of pro-caspase-9 leads to the cleavage and activation of pro-caspase-3 and pro-caspase-7, and the active forms of these latter two enzymes then execute apoptosis by cleaving numerous substrates (Wang and Youle, 2009).

Mitochondria are highly dynamic organelles that continually change their morphology by fission and fusion processes, which contributes to mitochondrial quality control and induction of apoptosis (Detmer and Chan, 2007; Suen et al., 2008). Mitochondrial fission and fusion are mediated by evolutionarily conserved members of the dynamin family of proteins. Fission is thus mediated by cytosolic dynamins such as Drp1 (dynamin-related protein 1) and Dyn2, whereas fusion is mediated by the membrane-anchored dynamins Mfn1-Mfn2 and Opa1, respectively, in mammals (Detmer and Chan, 2007; Lee et al., 2016). The fission process mediated by Drp1 appears to play a central role in the induction of cytochrome *c* release and subsequent apoptosis in various physiological and pathological contexts (Westermann, 2010).

Recent studies have shown that interactions with other organelles contribute to the regulation of mitochondrial dynamics. Sites of contact between mitochondria with ER (known as mitochondrion-associated membranes, MAMs) play an important role in the regulation of mitochondrial fission (Friedman et al., 2011) and have been implicated in that of apoptosis (Hoppins and Nunnari, 2012; Prudent et al., 2015; Yang et al., 2018). Lysosomes also participate in the regulation of mitochondrial dynamics (Wong et al., 2018). Mitochondria and peroxisomes share key regulators of their fission including Drp1 as well as Fis1 and Mff1 (Camões et al., 2009; Delille et al., 2009; Kobayashi et al., 2007; Schrader, 2006). However, whether peroxisomes also regulate mitochondrial dynamics and mitochondrion-mediated apoptosis has remained unclear.

We have now investigated the role of peroxisomes in the regulation of mitochondrial dynamics, caspase activation, and apoptosis by deleting *Pex3* or *Pex5* in mouse embryonic fibroblasts (MEFs) under conditions in which the cytosolic ROS level does not increase substantially. We found that deletion of either *Pex3* or *Pex5* resulted in fragmentation of mitochondria, the appearance of cytochrome *c* in the cytosol, and an increase in the amounts of cleaved caspase-9 and caspase-3. Importantly, restoration of Pex3 or Pex5 expression in the corresponding knockout (KO) MEFs attenuated these effects. Furthermore, we found that ablation of Pex3 greatly enhanced the induction of apoptosis by the DNA-damaging agent etoposide. Our results thus suggest that peroxisomes regulate mitochondrial dynamics, caspase activity, and cell death so as to reduce cellular sensitivity to damaging insults.

## Results

### Induction of mitochondrial fragmentation by *Pex3* deletion

To examine an acute effect of peroxisome deficiency, we took advantage of MEFs derived from *Pex3*^fl/fl^;*Rosa-Cre-ER*^T2^ mice, which are homozygous for a floxed allele of *Pex3* and harbor a tamoxifen-inducible transgene for Cre recombinase (Fig. S1). We immortalized these cells by introducing SV40 large T antigen and deleted *Pex3* by adding 4-hydroxytamoxifen. Immunoblot analysis detected Pex3 protein in control MEFs (not exposed to 4-hydroxytamoxifen) but not in Pex3 KO MEFs (Fig. 1 A). Immunofluorescence analysis also detected almost no punctate signals for Pex14 or for EGFP tagged with peroxisome-targeting signal 1 (PTS1) in the Pex3 KO MEFs (Fig. 1 B), indicating the successful depletion of peroxisomes in these cells.

**Figure 1.**
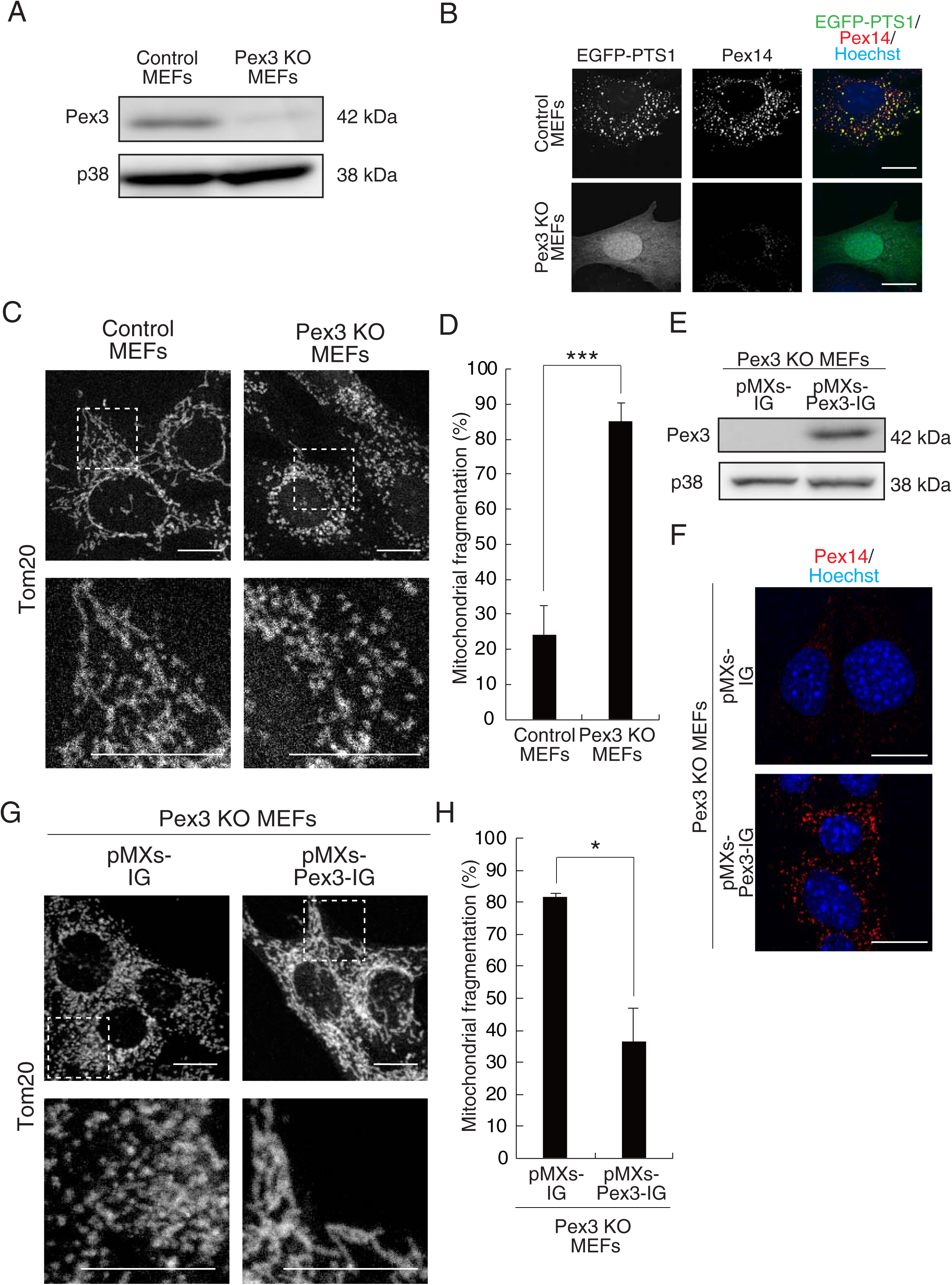
Deletion of *Pex3* induces mitochondrial fragmentation. **(A)** Immunoblot analysis of control and Pex3 KO MEFs with antibodies to Pex3 and to p38 (loading control). Data are representative of three independent experiments. **(B)** Immunofluorescence staining of control and Pex3 KO MEFs expressing EGFP-PTS1 with antibodies to Pex14. Nuclei were stained with Hoechst 33342. Scale bars, 20 μm. Data are representative of three independent experiments. **(C)** Immunofluorescence staining of control and Pex3 KO MEFs with antibodies to Tom20. The boxed regions in the upper panels are shown at higher magnification in the lower panels. Scale bars, 20 μm. **(D)** Quantification of mitochondrial fragmentation in control and Pex3 KO MEFs determined from images as in (C). Data are means ± SEM from three independent experiments. ***P < 0.005 (unpaired Student’s *t* test). **(E)** Immunoblot analysis of Pex3 in Pex3 KO MEFs infected with retroviruses encoding GFP either alone (pMXs-IG) or together with Pex3 (pMXs-Pex3-IG). Data are representative of three independent experiments. **(F)** Cells as in (E) were subjected to immunofluorescence staining with antibodies to Pex14. Nuclei were stained with Hoechst 33342. Scale bars, 20 μm. Data are representative of three independent experiments. **(G)** Cells as in (E) were subjected to immunofluorescence staining with antibodies to Tom20. The boxed regions in the upper panels are shown at higher magnification in the lower panels. Scale bars, 20 μm. **(H)** Quantification of mitochondrial fragmentation in images similar to those in (G). Data are means ± SEM from three independent experiments. *P < 0.05 (unpaired Student’s *t* test).

With the use of our Pex3 KO MEFs, we then set out to identify mitochondrial phenotypes of peroxisome deficiency that could be rescued by reintroduction of peroxisomes. We examined mitochondrial morphology by observing the intracellular distribution of Tom20, a mitochondrial outer membrane protein, and ATP synthase β, a mitochondrial inner membrane protein, and found that the extent of mitochondrial fragmentation was increased in Pex3 KO MEFs (Fig. 1,C and D, Fig. S2 A). Huygens-based quantification indicated that the size of mitochondria was significantly smaller in Pex3-deficient MEFs than that in control MEFs (Fig. S2 B). The length of mitochondria also became shorter in Pex3-deficient MEFs compared to that in control MEFs (Fig. S2 C). We then asked whether restoration of Pex3 expression in these Pex3-deficient cells would rescue this mitochondrial phenotype. Infection of Pex3 KO MEFs with a retrovirus encoding Pex3 increased the abundance of Pex3 protein and induced the formation of peroxisomes (Fig. 1,E and F). Importantly, this reintroduction of Pex3 resulted in elongation of mitochondria in the Pex3-deficient MEFs (Fig. 1,G and H), indicating that Pex3 suppresses mitochondrial fragmentation in a reversible manner.

### Induction of mitochondrial fragmentation by *Pex5* deletion

Given that the effect of *Pex3* deletion on mitochondrial morphology might have been the result of a Pex3-specific function unrelated to peroxisome formation, we examined whether deletion of a different Pex gene, *Pex5*, might confer a similar phenotype. Deletion of *Pex5* would be expected to result in loss of peroxisomal matrix proteins but retention of the peroxisomal membrane, whereas that of *Pex3* results in the complete loss of peroxisomes. To disrupt *Pex5* in MEFs, we adopted the CRISPR-Cas9 system with a gRNA targeted to *Pex5* (Fig. S3 A). We confirmed disruption of *Pex5* gene (Fig. S3 B) as well as the loss of Pex5 protein (Fig. 2 A) in the targeted cells. Examination of mitochondrial morphology revealed that the extent of mitochondrial fragmentation was increased in the Pex5-deficient MEFs compared with control (WT) MEFs (Fig. 2,B and C). Furthermore, this phenotype of Pex5 deficiency was rescued by retrovirus-mediated restoration of Pex5 expression (Fig. 2, D–F). These results thus provided further support for the notion that peroxisomes suppress mitochondrial fragmentation.

**Figure 2.**
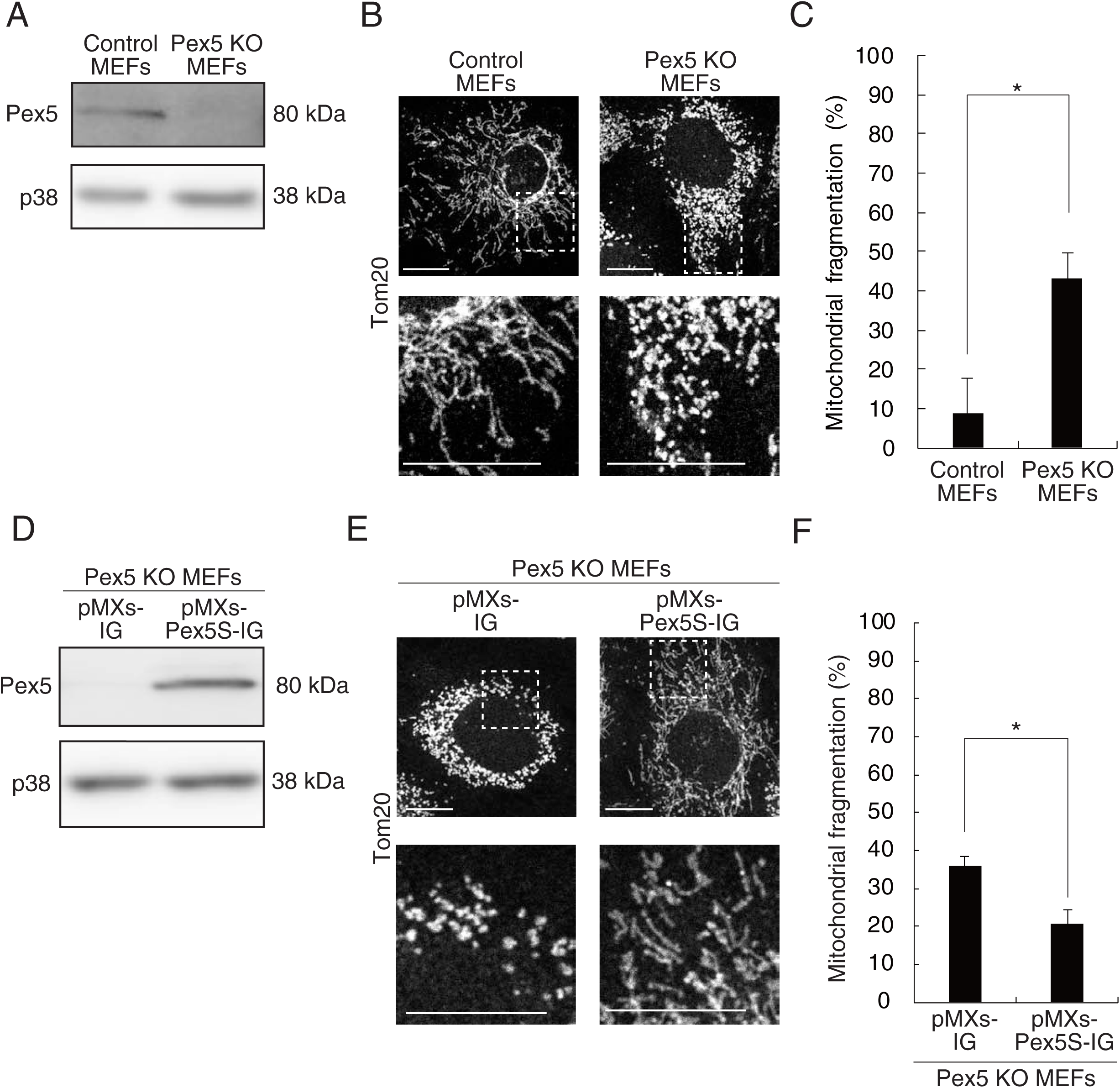
Deletion of *Pex5* induces mitochondrial fragmentation. **(A)** Immunoblot analysis of Pex5 in control and Pex5 KO MEFs. Data are representative of three independent experiments. **(B)** Immunofluorescence analysis of Tom20 in control and Pex5 KO MEFs. The boxed regions in the upper panels are shown at higher magnification in the lower panels. Scale bars, 20 μm. **(C)** Quantification of mitochondrial fragmentation in control and Pex5 KO MEFs determined from images as in (B). Data are means ± SEM from three independent experiments. *P < 0.05 (unpaired Student’s *t* test). **(D)** Immunoblot analysis of Pex5 in Pex5 KO MEFs infected with retroviruses encoding GFP either alone (pMXs-IG) or together with Pex5S (pMXs-Pex5S-IG). Data are representative of three independent experiments. **(E)** Immunofluorescence analysis of Tom20 in cells as in (D). The boxed regions in the upper panels are shown at higher magnification in the lower panels. Scale bars, 20 μm. **(F)** Quantification of mitochondrial fragmentation in control and Pex5 KO MEFs determined from images as in (E). Data are means ± SEM from four independent experiments. *P < 0.05 (unpaired Student’s *t* test).

### Induction of mitochondrial elongation by a peroxisome proliferator

We next examined whether an increase (rather than a decrease) in the number of peroxisomes might also affect mitochondrial morphology. We thus exposed MEFs to 4-phenylbutyrate (4-PBA), an inducer of peroxisome proliferation. We confirmed that 4-PBA increased the abundance of peroxisomes, as detected by immunostaining of Pex14, in control MEFs (Fig. 3 A). Furthermore, we found that 4-PBA induced mitochondrial elongation in these cells (Fig. 3, A, C, and D). Importantly, however, 4-PBA did not induce mitochondrial elongation or suppress mitochondrial fragmentation in Pex3 KO MEFs (Fig. 3, B, C, and E), suggesting that the induction of mitochondrial elongation by 4-PBA requires peroxisomes. Together, these results indicated that peroxisome abundance is a critical determinant of mitochondrial fission-fusion dynamics.

**Figure 3.**
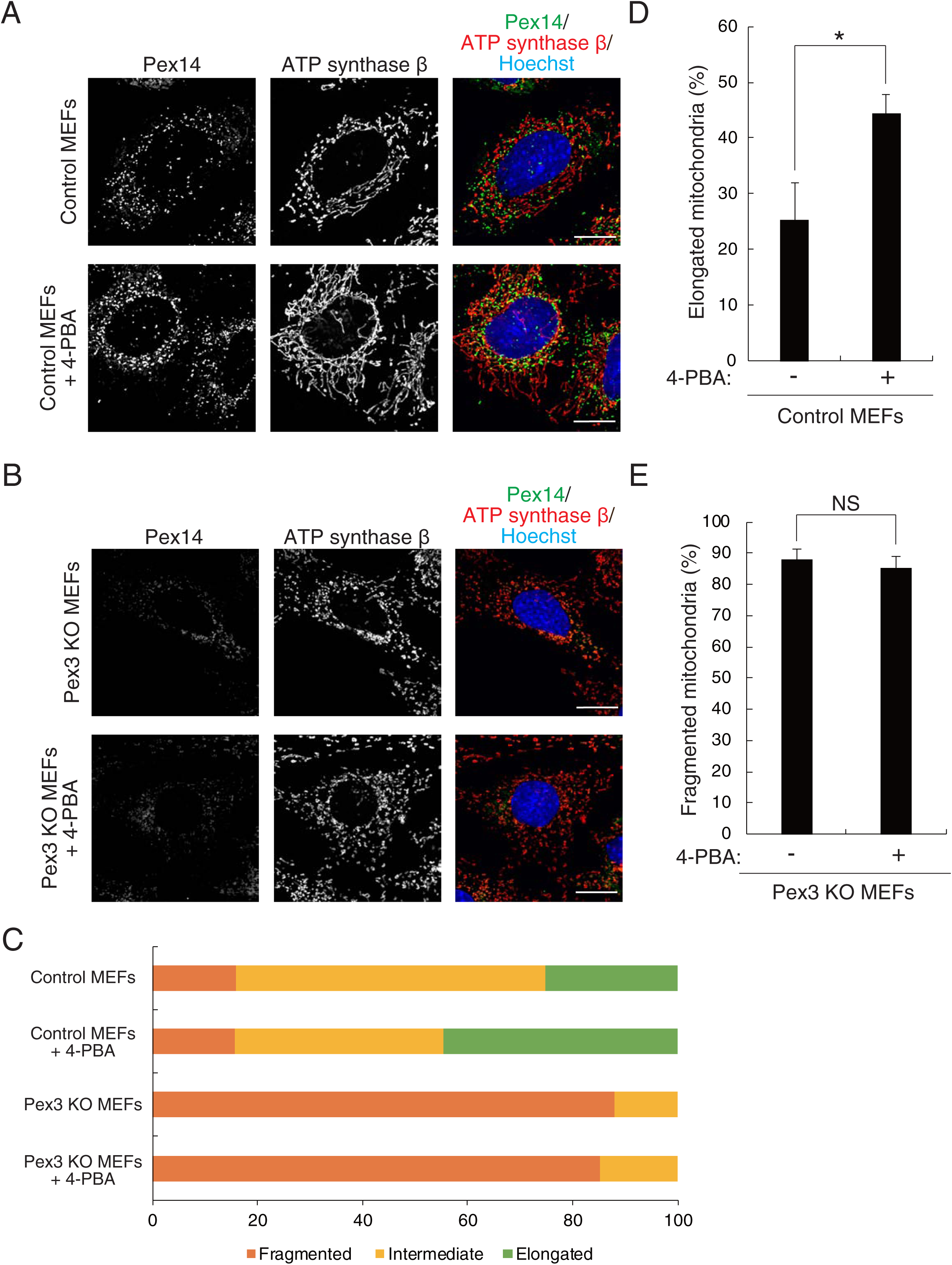
The peroxisome proliferator 4-PBA induces mitochondrial elongation. **(A)** Immunofluorescence staining of Pex14 and ATP synthase β (mitochondrial marker) in control MEFs cultured with or without 1 mM 4-PBA for 48 hours. Nuclei were stained with Hoechst 33342. Scale bars, 20 μm. **(B)** Quantification of mitochondrial elongation in control MEFs as in (A). Data are means ± SEM for four independent experiments. *P < 0.05 (unpaired Student’s *t* test). **(C)** Immunofluorescence analysis of Pex3 KO MEFs cultured and stained as in (A). **(D)** Quantification of mitochondrial fragmentation in Pex3 KO MEFs as in (C). Data are means ± SEM for four independent experiments. NS, unpaired Student’s *t* test. **(E)** Summary of the quantification of mitochondrial morphology in control and Pex3 KO MEFs cultured with or without 1 mM 4-PBA for 48 hours.

### Induction of cytochrome *c* diffusion by deletion of *Pex3*

Given the mitochondrial fragmentation apparent in peroxisome-deficient cells, we next examined mitochondrial structure in more detail by electron micrography (EM). Mitochondria were indeed smaller and shorter in Pex3 KO MEFs compared with control MEFs (Fig. 4, A–C), consistent with the results of confocal fluorescence microscopy (Fig. 1,C and D). The structure of cristae also appeared to have collapsed, with the presence of an indistinct and irregular inner membrane, in Pex3-deficient MEFs (Fig. 4 D), similar to the morphology previously observed in Pex5-deficient hepatocytes (Baumgart et al., 2001; Peeters et al., 2015).

**Figure 4.**
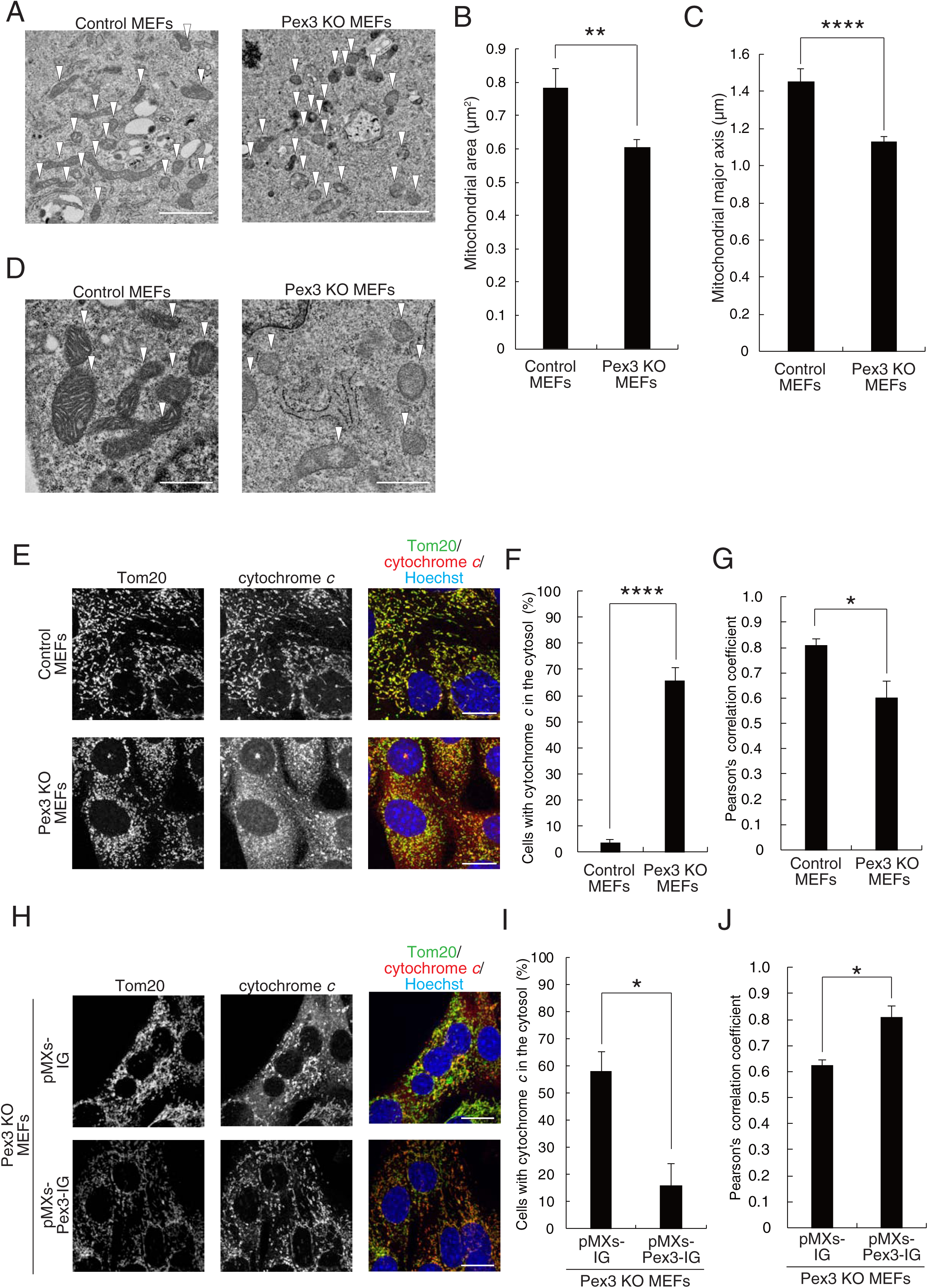
Deletion of *Pex3* induces cytochrome *c* diffusion. **(A)** EM of control and Pex3 KO MEFs. Scale bars, 1.0 μm. **(B and C)** Quantification of mitochondrial area and major axis, respectively, in images similar to those in (A). Data are means ± SEM for 21 cells of each genotype. **P < 0.01; ****P < 0.001 (unpaired Student’s *t* test). **(D)** High-magnification electron micrographs of control and Pex3 KO MEFs. Scale bars, 250 nm. **(E)** Immunofluorescence staining of Tom20 and cytochrome *c* in control and Pex3 KO MEFs. Nuclei were stained with Hoechst 33342. Scale bars, 20 μm. **(F)** Quantification of the cells with cytochrome *c* in the cytosol imaged as in (E). Data are means ± SEM from three independent experiments. ****P < 0.001 (unpaired Student’s *t* test). **(G)** Colocalization of cytochrome *c* with Tom20 as reflected by Pearson’s correlation coefficient (r) and determined from images as in (E). Data are means ± SEM from three independent experiments. *P < 0.05 (unpaired Student’s *t* test). **(H)** Immunofluorescence staining of Tom20 and cytochrome *c* in Pex3 KO MEFs infected with retroviruses encoding GFP either alone (pMXs-IG) or together with Pex3 (pMXs-Pex3-IG). Nuclei were stained with Hoechst 33342. Scale bars, 20 μm. **(I)** Quantification of the cells with cytochrome *c* in the cytosol imaged as in (H). Data are means ± SEM from three independent experiments. *P < 0.05 (unpaired Student’s *t* test). **(J)** Colocalization of cytochrome *c* with Tom20 as reflected by Pearson’s correlation coefficient and determined from images as in (H). Data are means ± SEM from three independent experiments. *P < 0.05 (unpaired Student’s *t* test).

Given that mitochondrial fragmentation and collapsed cristae are associated with the release of cytochrome *c* from these organelles and the intrinsic (mitochondrion-dependent) pathway of apoptosis (Otera et al., 2016; Suen et al., 2008), we examined whether *Pex3* deletion might affect the distribution of cytochrome *c*. Whereas cytochrome *c* immunoreactivity was detected almost exclusively in mitochondria of control MEFs, as shown by its overlap with that of Tom20, cytochrome *c* signals were diffusely distributed in the cytosol in addition to their punctate mitochondrial distribution in Pex3 KO MEFs (Fig. 4,E and F). A high cell density appeared to further increase the amount of cytochrome *c* in the cytosol of Pex3-deficient cells (Fig. S4). Quantitative analysis revealed that the colocalization of cytochrome *c* with Tom20, as reflected by Pearson’s correlation coefficient (r), was significantly reduced in Pex3 KO MEFs compared with control MEFs (Fig. 4 G), indicating that *Pex3* deletion indeed induced the diffusion of cytochrome *c*. Importantly, forced expression of Pex3 was sufficient to restore the normal (mitochondrial) distribution of cytochrome *c* in Pex3 KO MEFs (Fig. 4, H–J). Together, these results thus showed that Pex3 suppresses the diffusion of cytochrome *c*.

### Induction of cytochrome *c* diffusion by deletion of *Pex5*

We then examined whether deletion of *Pex5* results in a similar redistribution of cytochrome *c*. Cytochrome *c* signals were indeed found to be diffusely distributed in the cytosol of Pex5 KO MEFs (Fig. 5,A and B). We also confirmed that forced expression of Pex5 restored the mitochondrial localization of cytochrome *c* in the Pex5-deficient cells (Fig. 5,C and D). Together, our results thus indicated that functional peroxisomes, which require both Pex3 and Pex5, are necessary for suppression of cytochrome *c* diffusion.

**Figure 5.**
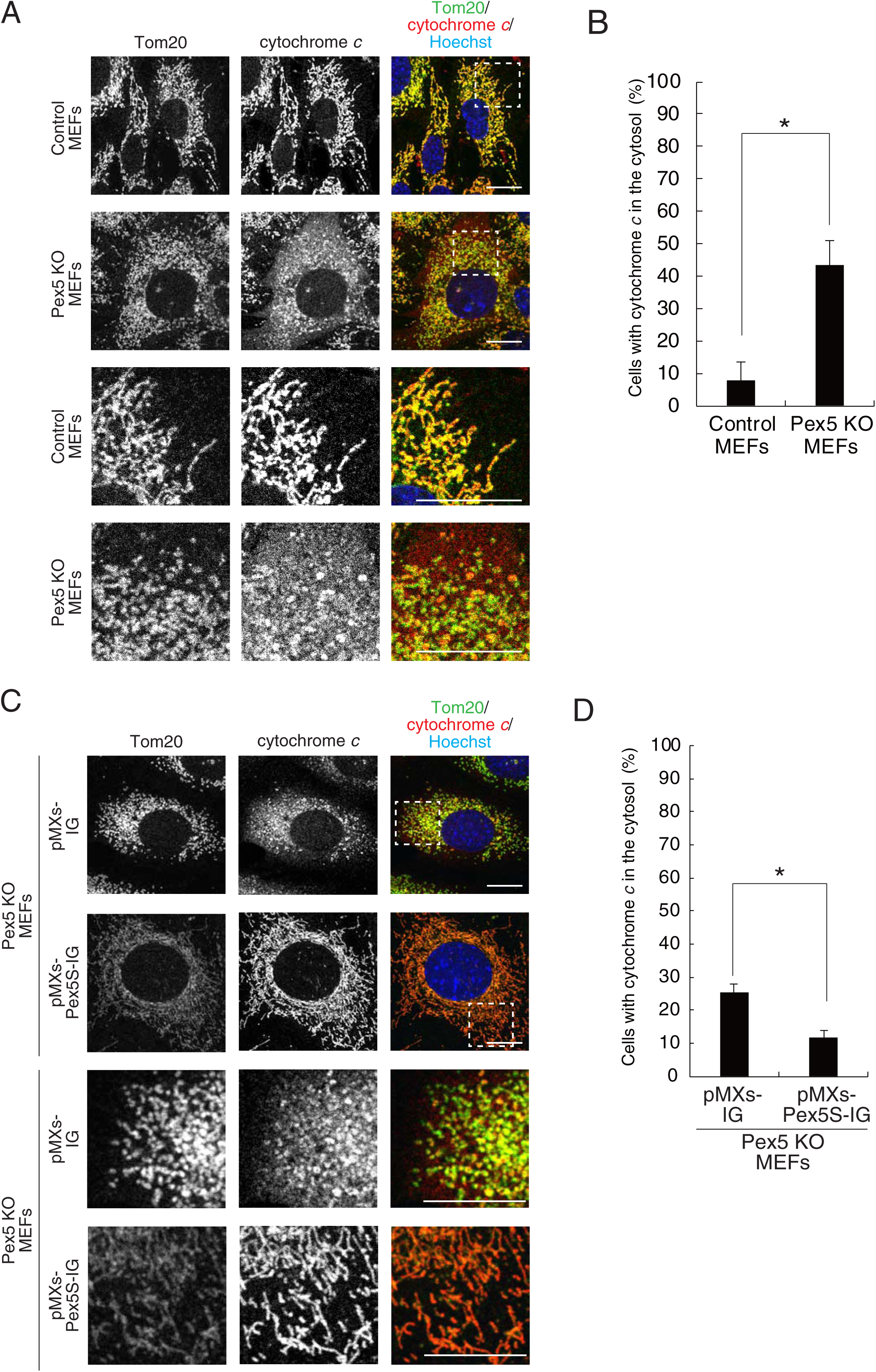
Deletion of *Pex5* induces cytochrome *c* diffusion. **(A)** Immunofluorescence staining of Tom20 and cytochrome *c* in control and Pex5 KO MEFs. Nuclei were stained with Hoechst 33342. The boxed regions in the upper panels are shown at higher magnification in the lower panels. Scale bars, 20 μm. **(B)** Quantification of the cells with cytochrome *c* in the cytosol imaged as in (A). Data are means ± SEM for three independent experiments. *P < 0.05 (unpaired Student’s *t* test). **(C)** Immunofluorescence staining of Tom20 and cytochrome *c* in Pex5 KO MEFs infected with retroviruses encoding GFP either alone (pMXs-IG) or together with Pex5S (pMXs-Pex5S-IG). Nuclei were stained with Hoechst 33342. The boxed regions in the upper panels are shown at higher magnification in the lower panels. Scale bars, 20 μm. **(D)** Quantification of the cells with cytochrome *c* in the cytosol imaged as in (C). Data are means ± SEM from four independent experiments. *P < 0.05 (unpaired Student’s *t* test).

### *Pex3* deletion without an overt change in ROS and respiration levels

We next addressed the mechanism by which the diffusion of cytochrome *c* is increased in peroxisome-deficient MEFs. Although previous studies have shown that long-term deletion of *Pex* genes results in ROS accumulation (Bülow et al. 2018, Rahim et al. 2016), the cytosolic ROS level of our Pex3-deficient MEFs as measured with CellROX did not appear to differ from that of control MEFs (Fig. 6,A and B). Under the same condition, the treatment of these MEFs with tert-butylhydroperoxide (TBHP), a ROS inducer, increased CellROX signals (Fig. 6,A and B). Moreover, control and Pex3-deficient MEFs did not show a detectable difference in mitochondrial ROS levels monitored by MitoSOX (Fig. 6,C and D). These results suggest that (relatively acute) *Pex3* deletion did not overtly increase ROS levels in mitochondria or the cytosol of our cultured MEFs. We also examined the rate of oxygen consumption in these cells, given that the abundance and activity of OXPHOS components are reduced after the deletion of *Pex* genes (Peeters et al., 2015). We again found, however, that control and Pex3-deficient MEFs did not differ significantly in their basal or ATP-linked rates of oxygen consumption (Fig. 6,E and F), suggesting that the mitochondrial OXPHOS system remains intact after *Pex3* deletion in MEFs. Considering that we did not observe overt ROS accumulation or altered oxygen consumption in these cells, it was unlikely that an increase in ROS levels was responsible for the induction of cytochrome *c* diffusion. Indeed, treatment of Pex3 KO MEFs with the ROS scavenger *N*-acetylcysteine (NAC) did not suppress mitochondrial fragmentation or cytochrome *c* diffusion, whereas such treatment did attenuate the TBHP-induced increase in CellROX signal intensity (Fig. S5).

**Figure 6.**
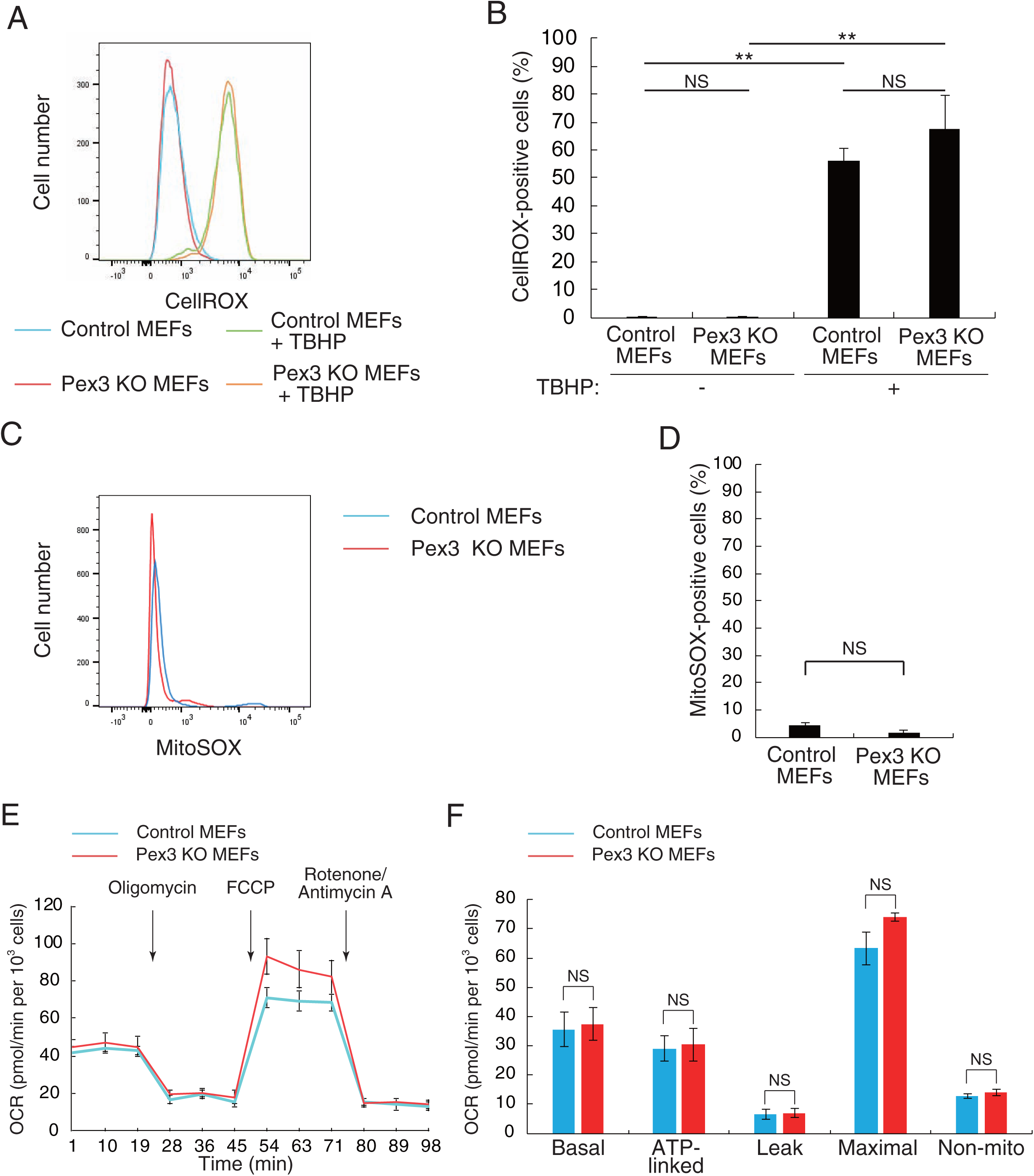
*Pex3* deletion without overt changes in ROS and respiration levels. **(A)** Representative flow cytometric analysis of cytosolic ROS levels as detected by CellROX staining in control and Pex3 KO MEFs incubated with or without 200 μM TBHP for 60 minutes. **(B)** Quantification of CellROX-positive cells as in (A). Data are means ± SEM from three independent experiments. **P < 0.01; NS, not significant (Scheffe’s test). **(C)** Representative flow cytometric analysis of mitochondrial ROS levels as detected by MitoSOX staining in control and Pex3 KO MEFs. **(D)** Quantification of MitoSOX-positive cells as in (C). Data are means ± SEM from six independent experiments. NS, not significant (unpaired Student’s *t* test). **(E)** Oxygen consumption rate (OCR) in control and Pex3 KO MEFs. Data are means ± SEM of triplicates from a representative experiment. **(F)** Basal, ATP-linked, proton-leak (+oligomycin), maximal (+CCCP), and nonmitochondrial (Non-mito, +rotenone/antimycin A) OCR determined as in (E). Data are means ± SEM from three independent experiments. NS, not significant (unpaired Student’s *t* test).

### Promotion of Drp1 association with mitochondria by deletion of *Pex3*

Given that Drp1 plays a pivotal role in mitochondrial fragmentation (fission) and cytochrome *c* release (Estaquier and Arnoult, 2007; Otera et al., 2016) and that Drp1 localizes not only to mitochondria but also to peroxisomes (Tanaka et al., 2006; Waterham et al., 2007), we examined whether *Pex3* deletion affects the abundance or subcellular localization of Drp1. Immunofluorescence analysis showed that the amounts of Drp1 both in the cytosol and associated with mitochondria appeared to increase in Pex3 KO MEFs compared with control MEFs (Fig. 7 A). Importantly, the extent of colocalization of Drp1 with Tom20 was significantly higher in Pex3-deficient MEFs than in control cells (Fig. 7 B), indicating that Pex3 ablation results in an increased localization of Drp1 to mitochondria.

To examine whether Drp1 is responsible for the mitochondrial fragmentation and cytochrome *c* diffusion observed in Pex3 KO MEFs, we suppressed the function of Drp1 by introducing a catalytically inactive mutant (K38A) of the protein that has been shown to act in a dominant negative manner (Frank et al., 2001). Expression of Drp1(K38A) indeed both restored the elongated morphology of mitochondria and attenuated cytochrome *c* diffusion in Pex3-deficient MEFs (Fig. 7 C–E). Together, these results thus suggested that *Pex3* deletion induces mitochondrial fragmentation and cytochrome *c* diffusion by promoting the localization of Drp1 to mitochondria.

**Figure 7.**
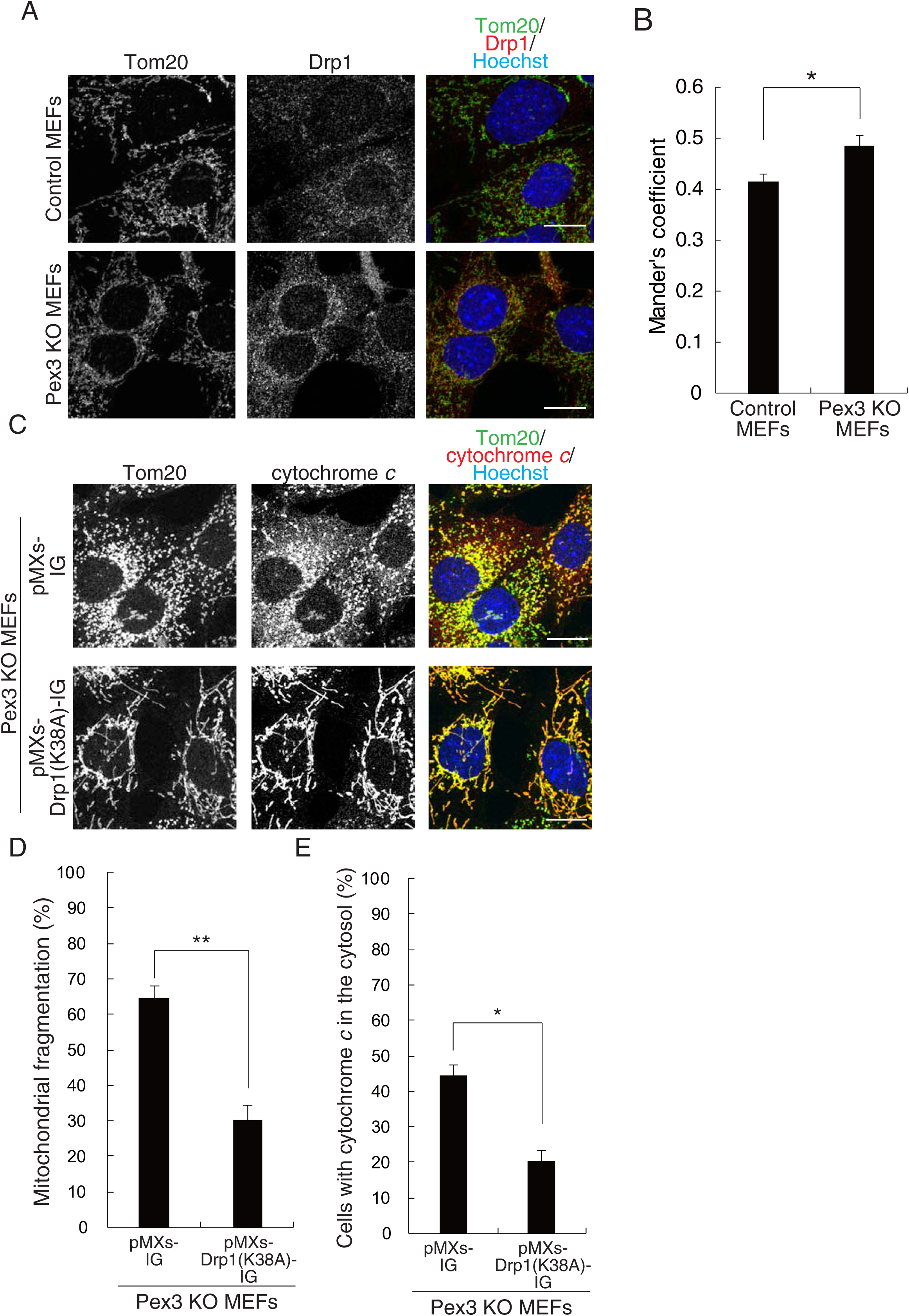
Deletion of *Pex3* promotes the association of Drp1 with mitochondria. **(A)** Immunofluorescence staining of Tom20 and Drp1 in control and Pex3 KO MEFs. Nuclei were stained with Hoechst 33342. Scale bars, 20 μm. **(B)** Colocalization of Drp1 with Tom20 as reflected by Manders’ M1 coefficient and determined from images as in (A). Data are means ± SEM for five independent experiments. *P < 0.05 (unpaired Student’s *t* test). **(C)** Immunofluorescence staining of Tom20 and cytochrome *c* in Pex3 KO MEFs infected with retroviruses encoding GFP either alone (pMXs-IG) or together with mutated Drp1 (Drp1(K38A)). Nuclei were stained with Hoechst 33342. Scale bars, 20 μm. Data are representative of three independent experiments. **(D)** Quantification of mitochondrial fragmentation in images similar to those in (C). Data are means ± SEM from three independent experiments. *P < 0.05 (unpaired Student’s *t* test). **(E)** Quantification of the cells with cytochrome *c* in the cytosol imaged as in (C). Data are means ± SEM from three independent experiments. *P < 0.05 (unpaired Student’s *t* test).

### Caspase activation and enhanced stress-induced apoptosis in Pex3-deficient cells

The release of cytochrome *c* from mitochondria triggers activation of the Apaf-1– caspase-9 complex (apoptosome) and caspase-3 and thereby induces apoptosis (Hyman and Yuan, 2012). We therefore examined the effect of *Pex3* deletion on caspase activity and found that the levels of the cleaved forms of caspase-9 and caspase-3 were increased in Pex3 KO MEFs compared with control MEFs (Fig. 8,A and B). These results thus suggested that Pex3 suppresses the activation of caspase-9 and caspase-3 under basal conditions in MEFs. In contrast, we did not detect any significant difference in the fraction of annexin V–positive (apoptotic) cells between control and Pex3-deficient MEFs (Fig. 8,C and D), suggesting that caspase activation induced by *Pex3* deletion is not sufficient to trigger apoptosis.

**Figure 8.**
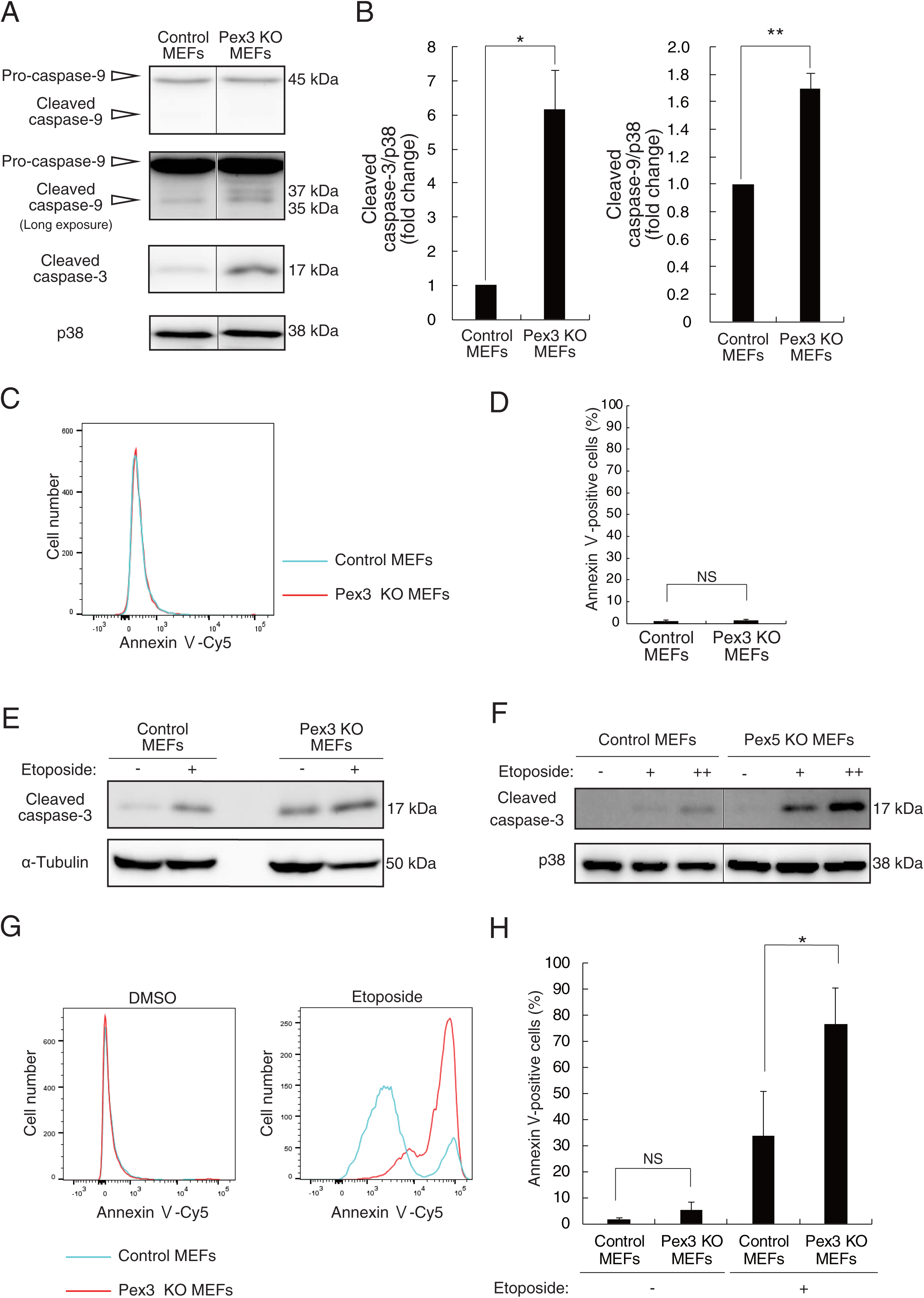
Deletion of *Pex3* induces caspase activation and enhances stress-induced apoptosis. **(A)** Immunoblot analysis of caspase-3 and caspase-9 in control and Pex3 KO MEFs. The pro and cleaved forms of the enzymes are indicated. Black vertical lines indicate noncontiguous lanes. **(B)** Quantification of the cleaved forms of caspase-3 and caspase-9 (normalized by p38) in blots similar to those in (A). Data are means ± SEM for three independent experiments. *P < 0.05, **P < 0.01 (unpaired Student’s *t* test). **(C)** Representative flow cytometric analysis of Cy5-labeled annexin V staining for control and Pex3 KO MEFs. **(D)** Quantification of cells positive for annexin V–Cy5 staining as in (C). Data are means ± SEM from four independent experiments. NS, not significant (unpaired Student’s *t* test). **(E and F)** Immunoblot analysis of the cleaved form of caspase-3 in control and either Pex3 KO (E) or Pex5 KO (F) MEFs that had been incubated in the absence (–) or presence of etoposide at 2 (+) or 4 (++) μM for 24 hours. Either α-tubulin or p38 was examined as a loading control. Black vertical lines indicate noncontiguous lanes. Data are representative of three independent experiments. **(G)** Representative flow cytometric analysis of annexin V–Cy5 staining for control and Pex3 KO MEFs that had been incubated for 24 hours with 2 μM etoposide or DMSO vehicle. **(H)** Quantification of cells positive for annexin V–Cy5 staining as in (G). Data are means ± SEM from three independent experiments. *P < 0.05; NS, not significant (unpaired Student’s *t* test).

We hypothesized that the elevated basal activity of caspases in Pex3 KO MEFs might increase the vulnerability of these cells to cellular stressors. To test this notion, we treated peroxisome-deficient cells with the DNA-damaging agent etoposide. Both Pex3-deficient MEFs and Pex5-deficient MEFs manifested increased levels of caspase-3 activation in response to etoposide treatment compared with the corresponding control MEFs (Fig. 8,E and F). Furthermore, annexin V staining revealed that Pex3 KO MEFs underwent apoptosis to a significantly greater extent than did control MEFs in response to etoposide treatment (Fig. 8,G and H). These results thus suggested that peroxisomes prevent excessive caspase activity and the induction of apoptosis, and that they thereby increase the resistance of cells to cellular stress such as that associated with DNA damage.

## Discussion

Although the cooperation between peroxisomes and mitochondria with regard to cellular metabolism has been extensively studied, the possible contribution of peroxisomes to mitochondrial fission-fusion dynamics has remained largely unknown. Our results now show that peroxisomes play a key role in determination of the balance between mitochondrial fission and fusion, with this balance being essential for a wide range of biological processes including cellular responsiveness to stressors (Detmer and Chan, 2007; Khacho et al., 2016; Weir et al., 2017). Indeed, our data also show that peroxisomes are important for protection of cells from mitochondrion-dependent apoptosis in response to DNA damage. Our study therefore provides a new basis for understanding the function of peroxisomes.

We found that the loss of peroxisomes induced the fragmentation of mitochondria, whereas some previous studies showed that the loss of peroxisomes promoted the enlargement of mitochondria (Bülow et al., 2018; Rahim et al., 2016). This difference may be attributable to the long-term ablation of peroxins in these previous studies, which likely resulted in secondary effects due to the accumulation of ROS and subsequent cellular damage. In the present study, we took advantage of a MEF culture in which the intracellular ROS level was not greatly increased after peroxin ablation and found that the mitochondrial fragmentation and cytochrome *c* diffusion induced by peroxin gene deletion were rescued by restoration of peroxin expression, indicating that these phenomena are primary effects of peroxisomal loss.

The mechanism responsible for peroxisome-mediated regulation of mitochondrial fission-fusion dynamics remains unknown. Given that the fission and fusion machineries of both organelles share components such as Drp1 and Fis1 in mammalian cells (Schrader, 2006; Tanaka et al., 2006; Waterham et al., 2007), peroxisomes and mitochondria may compete for these components. Indeed, we found that *Pex3* deletion increased Drp1 localization to mitochondria and that inhibition of Drp1 rescued mitochondrial fragmentation in Pex3 KO MEFs. These results thus implicate Drp1 function involved in the peroxisome-mediated regulation of mitochondrial dynamics. Several studies have also shown that peroxisomes are located adjacent to MAMs, which are described as mitochondrial constriction sites (Cohen et al., 2014; Friedman et al., 2011; Horner et al., 2011; Mattiazzi Usaj et al., 2015). Given that peroxisomes also make physical contact with mitochondria (Fransen et al., 2017), these observations raise the possibility that peroxisomes compete with ER for mitochondrial contact sites. Whether peroxisomes actually regulate MAM formation warrants future study. Furthermore, a recent study showed that lysosomes also make contacts with mitochondria and regulate mitochondrial fission (Wong et al., 2018). Indeed, >80% of mitochondrial fission sites were found to contact lysosomes, whereas <20% of such sites contacted peroxisomes. Peroxisome-mitochondrion contacts may thus hamper or promote the interaction between lysosomes and mitochondria, resulting in modulation of the mitochondrial fission process. Together, these previous and present observations reveal multiple types of interorganellar communication that coordinately regulate mitochondrial fission-fusion dynamics.

In this study, we propose that Drp1 mediates mitochondrial fragmentation and subsequent cytochrome *c* diffusion in peroxisome-deficient cells. However, it remains unknown what molecular mechanism underlies cytochrome *c* diffusion after Drp1-mediated mitochondrial fragmentation in peroxisome-deficient cells. One possibility is that peroxisomes compete with mitochondria for some components necessary for cytochrome *c* diffusion. Cytochrome *c* is known to be released through the pore composed of the Bcl-2 family members BAX and BAK at the outer membrane of mitochondria (Tait and Green, 2010). Intriguingly, Fujiki and colleagues reported that a fraction of BAK also localizes to peroxisomes (Hosoi et al., 2017). Elimination of peroxisomes may thus alter BAK’s localization from peroxisomes to mitochondria, and the increased mitochondrial BAK may thereby facilitate cytochrome *c* diffusion in peroxisome-deficient cells. It would be important to test the notion that molecules shared by mitochondria and peroxisomes mediate their interorganellar communications.

Our results revealed not only the fragmentation of mitochondria in response to the loss of peroxisomes, but also the elongation of mitochondria in response to treatment of cells with the peroxisome proliferator 4-PBA. These findings suggest that peroxisomal abundance is an important determinant of mitochondrial dynamics. Cellular conditions that affect the abundance of peroxisomes might thus also influence mitochondrial dynamics through peroxisomes. In this regard, cellular stressors such as UV light exposure and elevated ROS levels increase the number of peroxisomes in both plant and mammalian cells (Schrader and Fahimi, 2006). This increase in peroxisomal number or abundance may thus contribute to a protective response to allow cells to cope with stress via suppression of mitochondrial fragmentation and caspase activation. Such a notion is consistent with our present results showing that peroxisomes reduce cellular sensitivity to toxic insults.

Fatty acids, such as oleic acid, and a high-fat diet are also thought to increase the abundance of peroxisomes (Diano et al., 2011; Ishii et al., 1980; Lock et al., 1989; Reddy and Mannaerts, 1994; Veenhuis et al., 1987). It would thus be of interest to determine whether the high level of fatty acid synthesis apparent in adult neural stem-progenitor cells (Knobloch et al., 2013) confers resistance to cellular stress through an increase in the number of peroxisomes. Indeed, the abundance of peroxisomes is known to be high in radial glia cells preserved for a long period and to be reduced by aging (Ahlemeyer et al., 2007), with such changes possibly having consequences for mitochondrial regulation in these cells.

Mitochondrion-dependent activation of caspases contributes not only to removal of unnecessary cells during development or damaged cells exposed to stress stimuli but also to regulation of tissue stem cell differentiation and terminal differentiation of myoblasts, erythroblasts, and keratinocytes (Hollville and Deshmukh, 2017). Furthermore, nonapoptotic caspase activation plays a key regulatory role in the pruning of neurites and the formation and maturation of neural circuits in the nervous system (Unsain and Barker, 2015). For example, caspase-9 is necessary for axon pruning in dorsal root ganglion neurons and cervical sympathetic neurons (Cusack et al., 2013; Simon et al., 2012). Nonapoptotic caspase activation is also implicated in regulation of the internalization of AMPA-sensitive glutamate receptors, which contributes to long-term depression in hippocampal neurons (Li et al., 2010). Nonapoptotic activation of caspases is thus essential for the control of various cellular processes. The activation of caspases at a sublethal level in peroxisome-deficient cells observed in the present study suggests that peroxisomes limit caspase activation under low-stress conditions. It will be of interest to examine the possible role of peroxisomes in various biological processes that require nonapoptotic caspase activation.

Individuals with Zellweger syndrome and peroxin-deficient mice manifest severe defects in various organs including the brain, bone, muscle, kidney and liver. The mechanisms underlying this broad range of abnormalities remain unknown, however. Dysfunction of the mitochondrial fusion machinery also gives rise to neurodegenerative diseases, muscle atrophy, and osteogenic abnormalities (Chen et al., 2010; Detmer and Chan, 2007; Romanello et al., 2010; Touvier et al., 2015). Degeneration of Purkinje cells, one of the most prominent features of patients with Zellweger syndrome (Barry and O’Keeffe, 2013; Trompier et al., 2014), is thus also observed in mice with Purkinje cell–specific deficiency of Mfn2 (Chen et al., 2007). The peroxisome-dependent regulation of mitochondria uncovered in the present study therefore raises the possibility that excessive mitochondrial fragmentation plays a causal role in the pathogenesis of Zellweger syndrome. If so, our findings may provide a basis for the development of new therapies for this lethal disease.

## Materials and methods

### Immunoblot analysis

Immunoblot analysis was performed as described previously (Okazaki et al., 2013). Immune complexes were detected with a chemiluminescence reagent [100 mM Tris-HCl (pH 8.5), 1.25 mM luminol, 0.2 mM coumaric acid, 0.009% H2O2] and an Image Quant LAS4000 instrument (GE Healthcare). Blot intensities were measured with Image J software.

### Immunofluorescence microscopy

Cells were fixed with 4% formaldehyde for 10 minutes at 37°C, permeabilized with 0.2% Triton X-100 in PBS for 5 minutes, and incubated for 30 minutes in PBS containing 2% FBS and 2% BSA (blocking buffer). They were then exposed first for 24 hours at 4°C to primary antibodies in blocking buffer and then for 1 hour at room temperature to Alexa Fluor–conjugated secondary antibodies (Thermo Fisher Scientific) and Hoechst 33342 in blocking buffer. Moviol were used as mounting medium. Images were acquired with a TCS SP5 confocal microscope (Leica) and were processed with Photoshop CS software (Adobe). Pearson’s correlation coefficient (r) for the colocalization of Tom20 and cytochrome c as well as Manders’ M1 coefficient for the colocalization of Tom20 and Drp1 were calculated with Coloc 2 of Fiji.

### Morphological quantification of mitochondria

In Figure S2, samples were prepared in the same way as in the immunofluorescence experiments, except that ProLong Diamond were used as mounting medium. Images were deconvoluted in Huygens software (Scientific Volume Imaging). After the deconvolution process, voxels and length of mitochondria were calculated with object analyzer, Huygens. 35 and 30 images for Control and Pex3 KO MEFs respectively, from three independent experiment were analyzed in this quantification.

### EM

Cells were fixed in 0.1 M phosphate buffer (pH 7.2) containing 2% glutaraldehyde and 2% paraformaldehyde, exposed to 1% OsO4, dehydrated, and embedded in Epon 812. Ultrathin sections (60 nm) were cut with an ultramicrotome (UC6, Leica Microsystems), stained with uranyl acetate and lead citrate, and examined with a Hitachi HT7700 electron microscope. The area and major axis of mitochondria in images were measured with the use of Fiji software.

### Measurement of ROS

Cells were incubated with 5 μM MitoSOX or 500 nM CellROX for 30 minutes at 37°C, isolated by exposure to trypsin, and resuspended in PBS containing 3% FBS for analysis with a FACSAria flow cytometer (BD Biosciences).

### Measurement of oxygen consumption rate

The oxygen consumption rate (OCR) of cells was measured with the use of a Seahorse XF24 Extracellular Flux Analyzer (Seahorse Biosciences). Cells were plated in 24-well Seahorse plates and cultured overnight, after which the medium was replaced with Seahorse XF Base medium supplemented with 10 mM glucose, 1 mM pyruvate, and GlutaMAX (2 ml/liter, Thermo Fisher Scientific). The cells were placed in a 37°C incubator without CO_2_ before loading into the analyzer. After measurement of basal respiration, the cells were exposed to 1 μM oligomycin to measure the proton leak, to 1 μM carbonylcyanide m-chlorophenylhydrazone (CCCP) to measure the maximal OCR, and to 0.5 μM rotenone and 0.5 μM antimycin A to measure the nonmitochondrial OCR. The ATP-linked OCR was calculated by a subtraction proton leak from basal OCR. Cells plated simultaneously in 96-well plates were counted to normalize OCR values.

### Annexin V binding assay

Cells were stained with Cy5-coupled annexin V (Promokine) according to the manufacturer’s instructions. Flow cytometric analysis of the stained cells was performed with a FACSAria flow cytometer (BD Biosciences).

### Cell culture and transfection

MEFs and Plat-E cells (Morita et al., 2000) were maintained in DMEM supplemented with 10% FBS and 1% penicillin-streptomycin. Plat-E cells were transfected with the use of the GeneJuice Transfection Reagent (Merck Millipore), whereas transfection of MEFs was performed with Lipofectamine 2000 or with Lipofectamine and PLUS Reagents (Thermo Fisher Scientific).

### Deletion of *Pex3*

C57BL/6 mice harboring the *Pex3*^tm3a(EUCOMM)Wtsi^ allele obtained from the EUCOMM (European Conditional Mouse Mutagenesis Program) consortium were crossed with *Act-FLP* transgenic mice (Kono et al., 2017) to remove the FRT-flanked region and subsequently with *Rosa-CreER*^T2^ transgenic mice (obtained from the U.S. National Cancer Institute) (Fig. S1). Mice heterozygous for the floxed allele of *Pex3* were mated, and the resulting homozygous embryos were isolated for preparation of MEFs. The MEFs were immortalized by the introduction of SV40 large T antigen as described previously (Ando et al., 2000), and they were then treated with 1 nM 4-hydroxytamoxifen to remove the loxP-flanked region. Immortalized MEFs treated with ethanol vehicle instead of 4-hydroxytamoxifen were prepared as control cells.

### Deletion of *Pex5*

3T3 MEFs (kindly provided by H. Ichijo) were transfected with the KO vector (see Plasmids below) and were then sorted with a FACSAria flow cytometer (BD Biosciences) to obtain GFP-positive cells, which were seeded as single cells in a 96-well plate.

### Genomic PCR analysis

For confirmation of *Pex5* deletion in MEFs, the cells were collected and lysed with genotyping buffer [50 mM KCl, 10 mM Tris-HCl (pH 8.3), 1.5 mM MgCl2, gelatin (0.1 mg/ml), 0.45 % NP-40, 0.45% Tween 20, proteinase K (500 μg/ml, Kanto Chemical)] or lysis buffer [1% SDS, 10 mM EDTA, 50 mM Tris-HCl (pH 8.1)] and were then incubated consecutively at 55°C for 3 hours and 98°C for 10 minutes. The *Pex5* locus was amplified by PCR with the use of KOD FX Neo (Toyobo) and the forward and reverse primers 5′-TCCCTTCCCCCAGCCCACTCCGGGTGCCTC-3′ and 5′-TCGGCGATGAATTCTTGGGACCAGTCGGTCTCATT-3′, respectively. The PCR products were ligated into PCR-Blunt (Thermo Fisher Scientific) for sequencing by Eurofin Genomics.

### Retrovirus-mediated expression of Pex3 or Pex5, Drp1(K38A)

Plat-E cells were transfected with pMXs-IG or either pMXs-Pex3-IG vector encoding human Pex3 or pMXs-Pex5S-IG vectors encoding Chinese hamster Pex5S, or pMXs-Drp1(K38A)-IG vector encoding rat Drp1(K38A) (Morita et al., 2000). After three days, the culture supernatants were harvested for isolation of retroviruses. Pex3 KO or Pex5 KO MEFs were infected with the corresponding peroxin retrovirus or the control virus, after which the cells were sorted with a FACSAria flow cytometer (BD Biosciences) to obtain GFP-positive cells. For preparing Drp1(K38A) infected cells, Pex3 KO MEFs were infected with the Drp1(K38A) retrovirus or the control virus and infected cells were sorted in the same way as above.

### Plasmids

The plasmid pUcD2Hyg/EGFP-PTS1 was described previously (Tamura et al., 1998). Full-length cDNAs for human Pex3 or Chinese hamster Pex5S (His-ClPex5S-HA) (Ghaedi et al., 2000; Matsumura et al., 2000) were subcloned into the BamHI and XhoI sites of the pMXs-IG vector (kindly provided by T. Kitamura). The p3xFLAG-ratDrp1K38A plasmid encoding rat Drp1(K38A) was kindly provided by N. Ishihara. Full-length cDNAs for rat Drp1(K38A) were subcloned into the EcoRI and XhoI sites of the pMXs-IG vector. For generation of the CRISPR vector for *Pex5* deletion, a pair of oligonucleotides encoding the gRNA (forward, 5′-CACCGCTGGTCACCATGGCAATGC-3′; reverse, 5′-AAACGCATTGCCATGGTGACCAGC-3′) was annealed and ligated into the px458 vector (Ran et al., 2013).

### Reagents

NAC was obtained from Sigma-Aldrich. Etoposide, Hoechst 33342, a CellROX Green Flow Cytometry Assay Kit (including TBHP, NAC), ProLong Diamond, and MitoSOX Red Reagents were obtained from Thermo Fisher Scientific. Seahorse XF Cell Mito Stress Test Kit were obtained from Primetech. 4-PBA was obtained from Tocris.

### Antibodies

Polyclonal and monoclonal antibodies to cleaved caspase-3 were obtained from Cell Signaling; antibodies to p38 and to Tom20 were from Santa Cruz Biotechnology; those to α-tubulin were from Sigma; those to cytochrome c and to Drp1 were from BD Pharmingen; those to Pex14 were from Proteintech; those to Pex3 were from Atlas Antibodies; those to ATP synthase β were from Thermo Fisher Scientific; those to caspase-9 were from MBL Life Science; and those to Pex5 were described previously (Okumoto et al., 2014).

### Statistical analysis

Quantitative data are presented as means ± SEM and were compared with Scheffe’s test or the unpaired Student’s *t* test. A P value of <0.05 was considered statistically significant.

## Acknowledgments

We thank M. Okajima (Graduate School of Pharmaceutical Sciences, The University of Tokyo) for technical assistance, N. Ishihara (Institute of Life Science, Kurume University) and T. Kitamura (The institute of Medical Sciences, The University of Tokyo) for providing the plasmids, H. Ichijo (Graduate School of Pharmaceutical Sciences, The University of Tokyo) for providing the cells. We also appreciate our laboratory members for discussions.

---

**Supplemental Figure S1.**
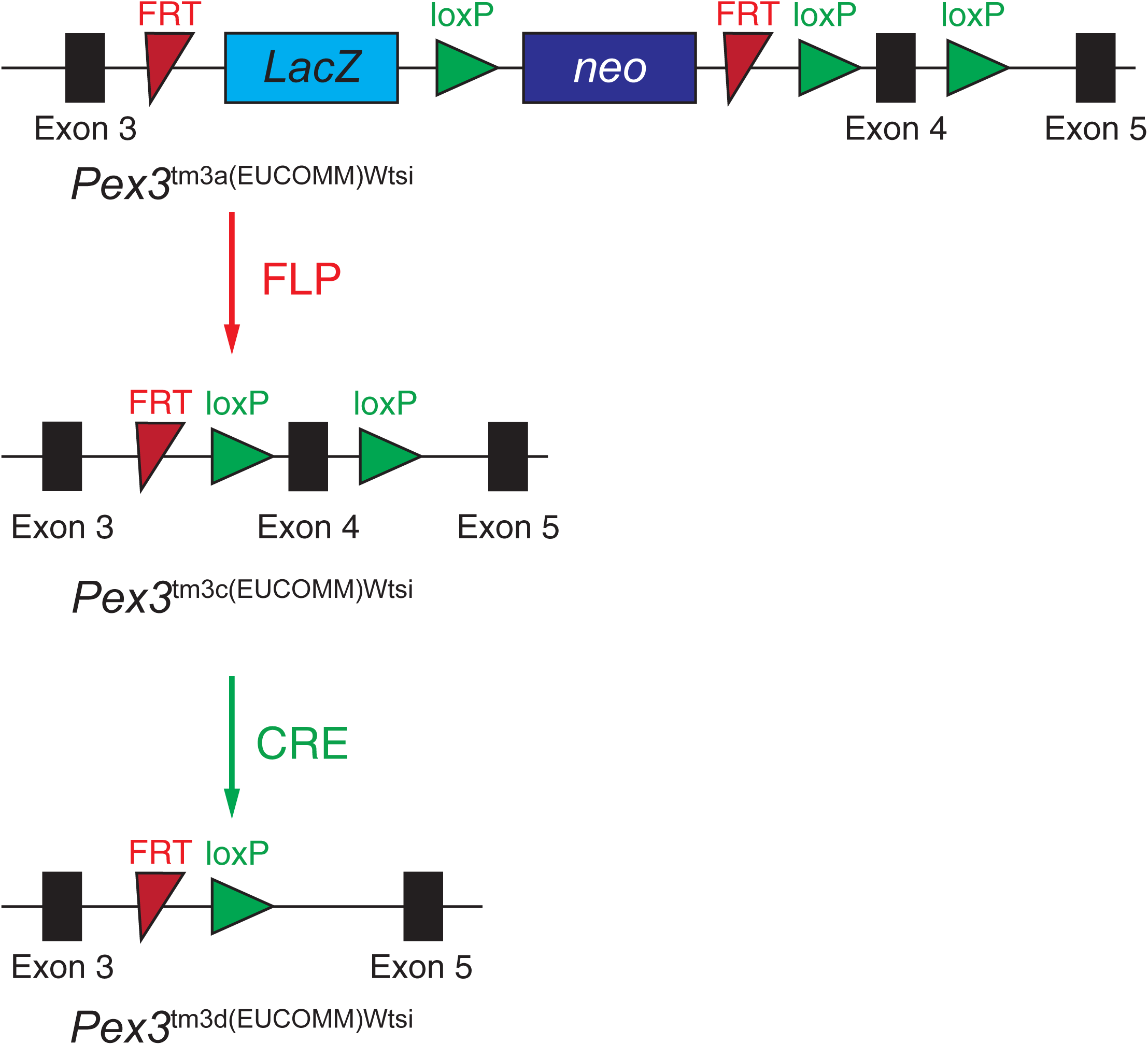
Scheme for *Pex3* disruption. The region of the *Pex3* locus spanning exons 3 to 5 for *Pex3*^tm3a(EUCOMM)Wtsi^, *Pex3*^tm3c(EUCOMM)Wtsi^, and *Pex3*^tm3d(EUCOMM)Wtsi^ alleles. See Materials and methods for details.

**Supplemental Figure S2.**
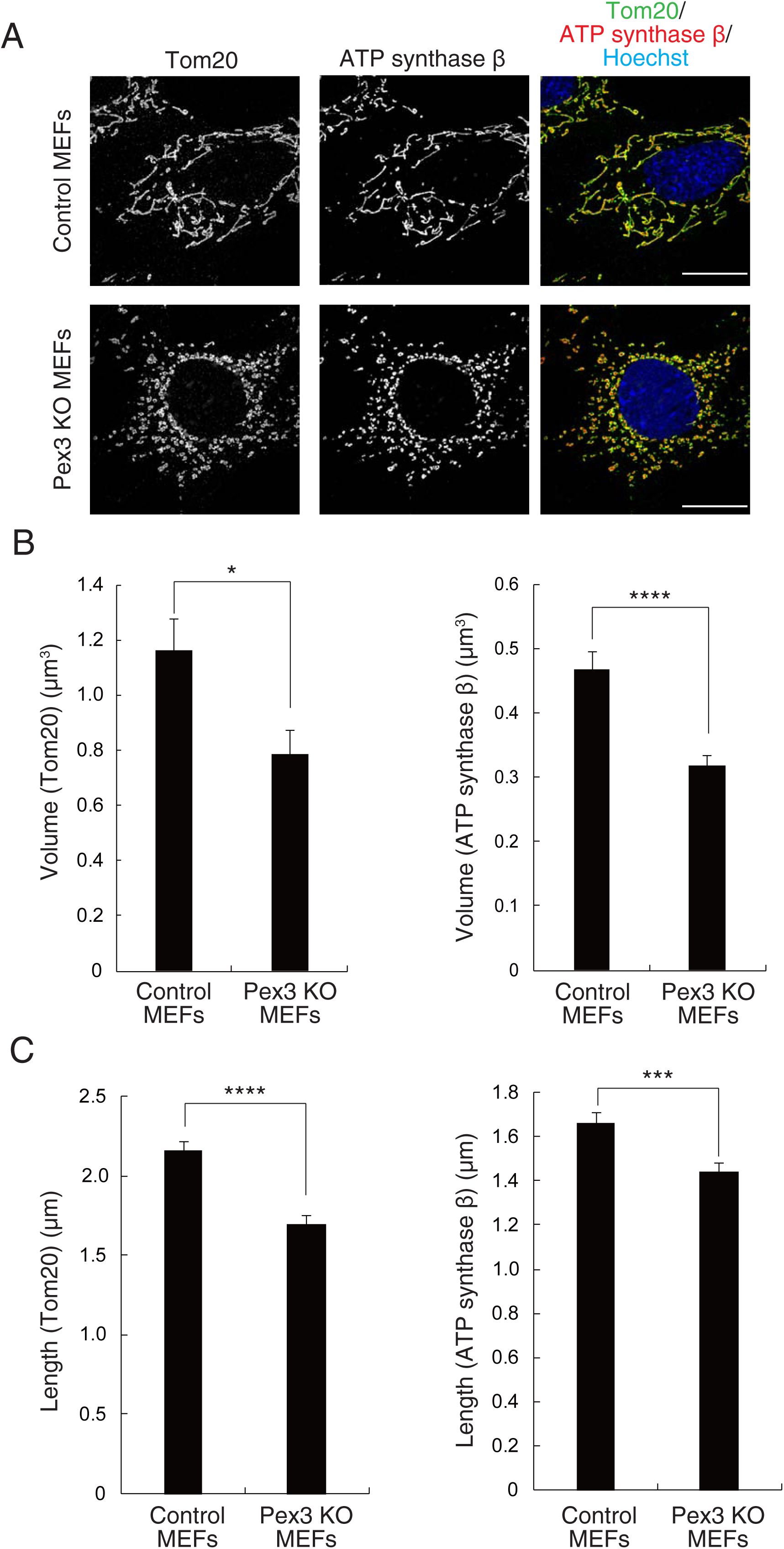
Deletion of *Pex3* induces mitochondrial fragmentation. **(A)** Deconvoluted Immunofluorescence images of control and Pex3 KO MEFs with antibodies to Tom20 and ATP synthase β. Scale bars, 20 μm. **(B)** Quantification of mitochondrial volume with object analyzer, Huygens and determined from imaged as in (A). Data are means ± SEM from 35 cells in control MEFs and 30 cells in Pex3 KO MEFs from three independent experiments. *P < 0.05; ****P < 0.001 (unpaired Student’s *t* test). **(C)** Quantification of mitochondrial length with object analyzer, Huygens and determined from imaged as in (A). Data are means ± SEM from 35 cells in control MEFs and 30 cells in Pex3 KO MEFs from three independent experiments. ***P < 0.005; ****P < 0.001 (unpaired Student’s *t* test).

**Supplemental Figure S3.**
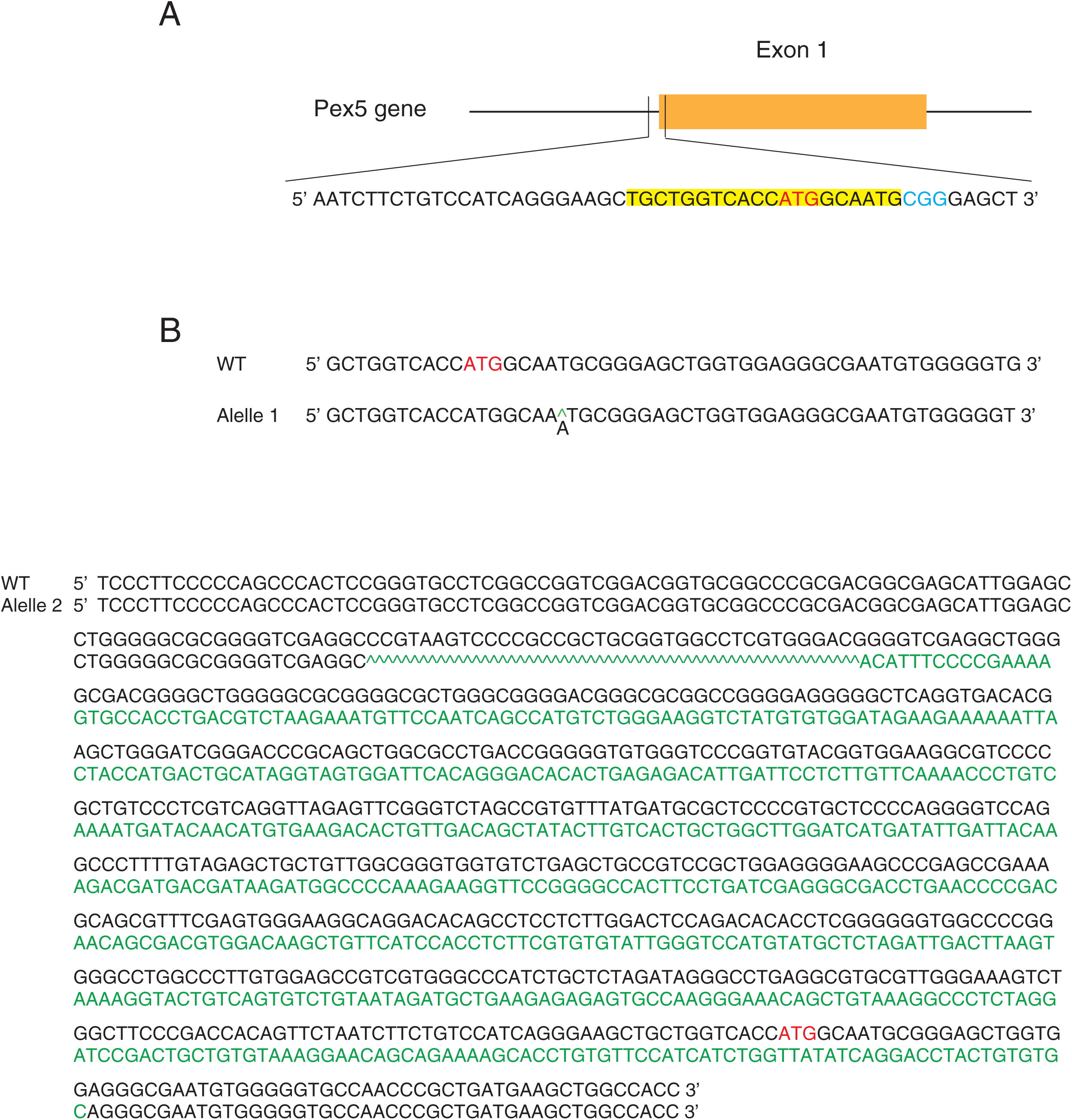
Indel of *Pex5* induced with the CRISPR-Cas9 system. **(A)** Schematic representation of the targeting *Pex5* with a gRNA. The gRNA sequence is shown in yellow, protospacer adjacent motif (PAM) in blue, and start codon in red. **(B)** Sequencing of *Pex5* genomic DNA from Pex5 KO MEFs. Insertion or replacement of nucleotides is shown in green. The start codon is presented in red.

**Supplemental Figure S4.**
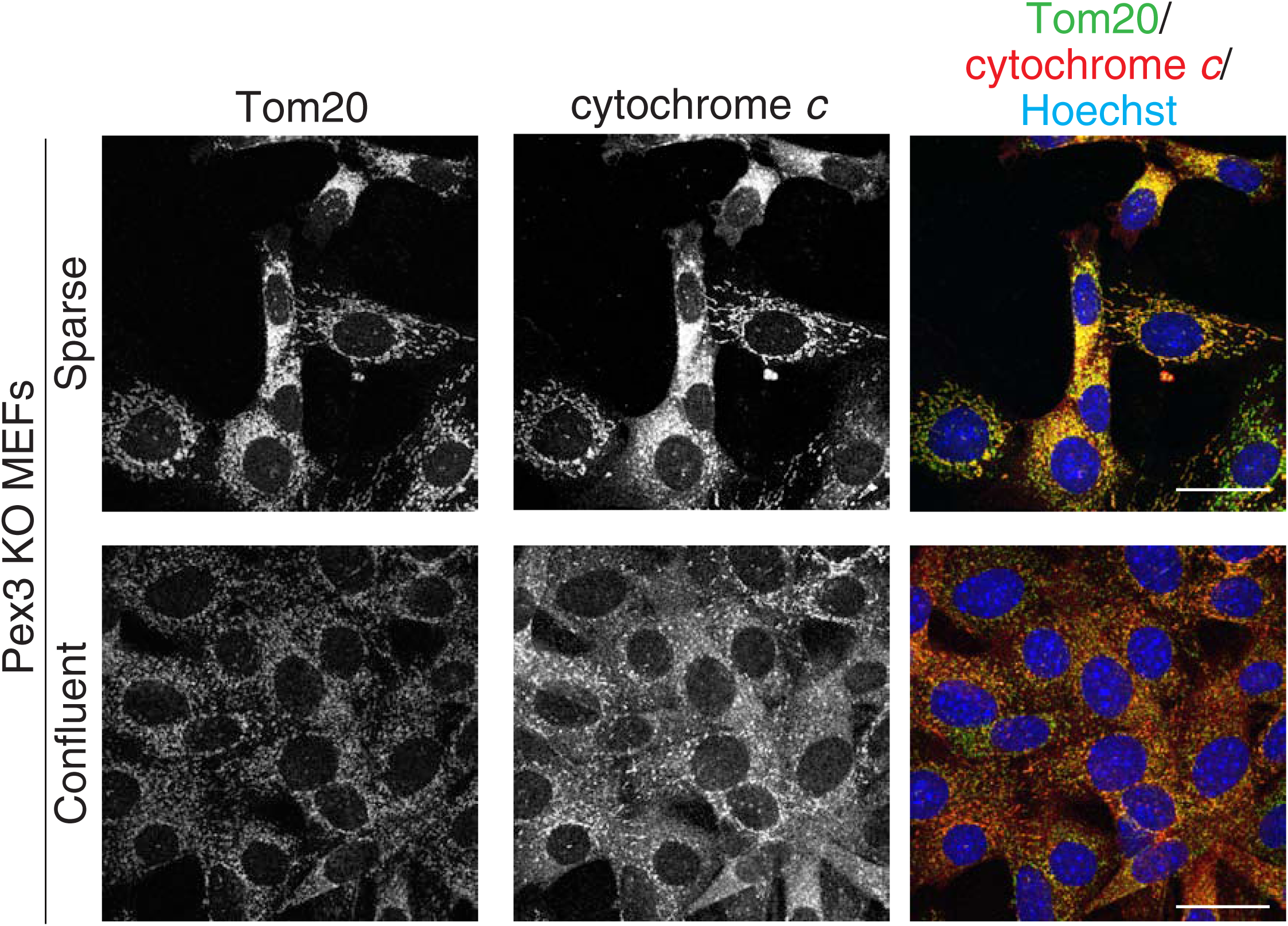
A high cell density enhances cytochrome *c* diffusion of Pex3 KO MEFs. Pex3 KO MEFs seeded at low or high cell densities were subjected to immunofluorescence staining of Tom20 and cytochrome *c*. Nuclei were stained with Hoechst 33342. Data are representative of three independent experiments. Scale bars, 40 μm.

**Supplemental Figure S5.**
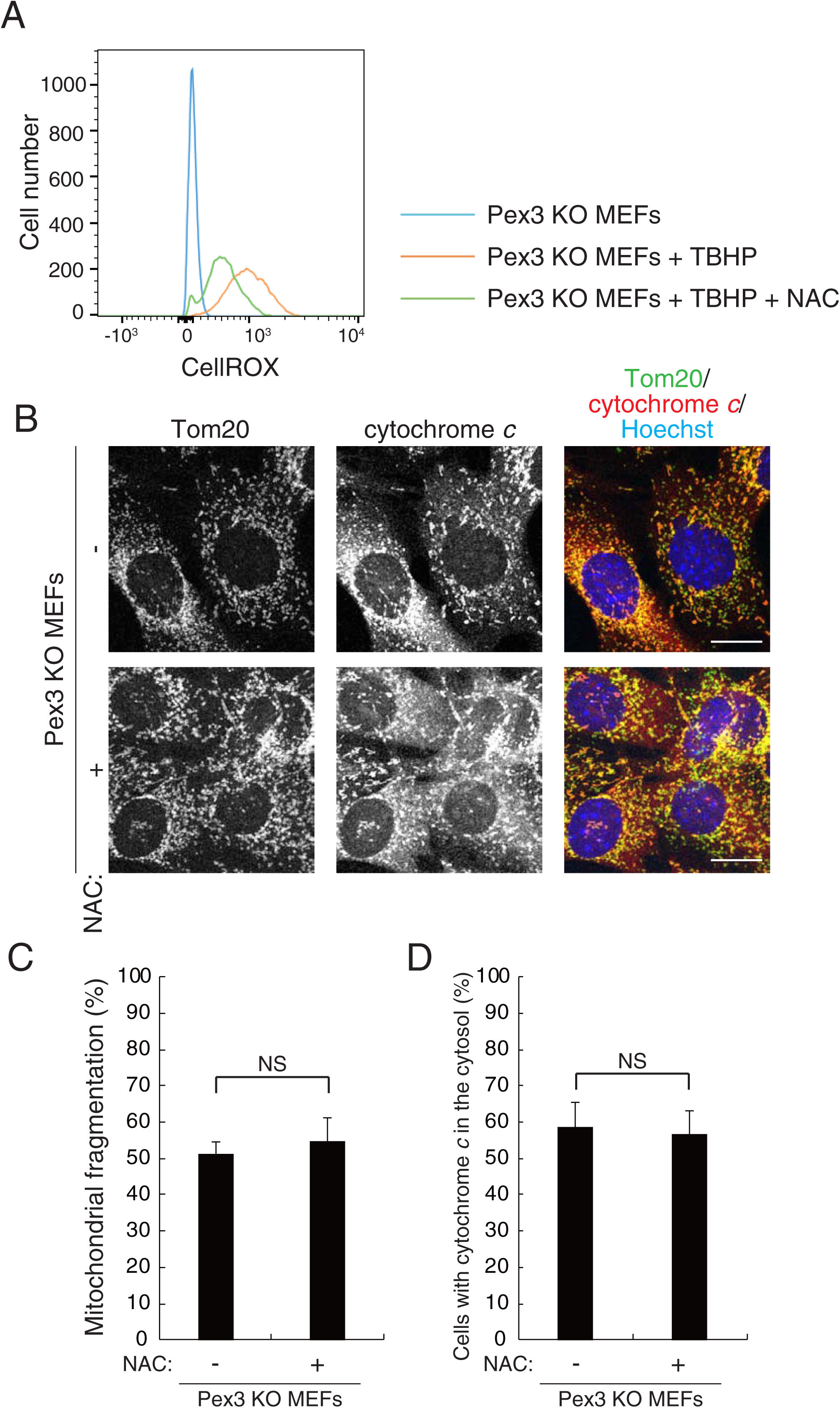
Mitochondrial fragmentation and cytochrome *c* diffusion are not mediated by ROS in Pex3 KO MEFs. **(A)** Flow cytometric analysis of cytosolic ROS as detected by CellROX staining in Pex3 KO MEFs incubated with or without 5 mM NAC for 1 hour and then in the additional absence or presence of 200 μM TBHP for 1 hour. Data are representative of three independent experiments. **(B)** Immunofluorescence staining of Tom20 and cytochrome *c* in Pex3 KO MEFs incubated with or without 5 mM NAC for 6 hours. Nuclei were also stained with Hoechst 33342. Scale bars, 20 μm. **(C and D)** Quantification of mitochondrial fragmentation and the diffusion of cytochrome *c* into the cytosol, respectively, for images similar to those in (B). Data are means ± SEM for three independent experiments. NS, not significant (unpaired Student’s *t* test).

## Nonstandard abbreviations

Drp1: dynamin-related protein 1
CCCP: carbonylcyanide m-chlorophenylhydrazone
KO: knockout
MAM: mitochondrion-associated membrane
MEF: mouse embryonic fibroblast
NAC: *N*-acetylcysteine
OXPHOS: oxidative phosphorylation
4-PBA: 4-phenylbutyrate
Pex: peroxin
PTS1: peroxisome-targeting signal 1
ROS: reactive oxygen species
TBHP: *tert*-butylhydroperoxide

